# Beneficial effects of *Spirogyra Neglecta Extract* on antioxidant and anti-inflammatory factors in streptozotocin-induced diabetic rats

**DOI:** 10.1101/485219

**Authors:** Behzad Mesbahzadeh, Seyed Ali Rajaei, Parnia Tarahomi, Seyed Ali Seyedinia, Mehrnoush Rahmani, Fatemeh Rezamohamadi, Muhammad Azam Kakar, Nasroallah Moradi-kor

## Abstract

**Objectives:** This study was conducted to evaluate the effects of oral supplementation of *Spirogyra* algae on oxidative damages and inflammatory responses in streptozotocin (STZ)-induced diabetic rats.

**Methods:** Diabetes was induced by administration of 55 mg/kg of streptozotocin. A total of sixty-four rats were divided into eight groups of eight rats each as follows:1) non-diabetic control; 2, 3, and 4) non-diabetic rats treated with 15, 30, and 45 mg of *Spirogyra* algae/kg/d; 5) control diabetic; and 6, 7, and 8) diabetic rats treated with 15, 30, and 45 mg of *Spirogyra* algae extract. At the end of the trial, the serum concentrations of glucose, interleukin-6 (IL-6), tumor necrosis factor-α (TNF-α), malondialdehyde (MDA), glutathione (GSH), total antioxidant status (TAS), C-reactive protein (CRP), insulin, triglycerides, and cholesterol were examined by specified procedures.

**Results:** Our findings indicated that the administration of STZ significantly increased the serum concentrations of glucose, triglycerides, cholesterol, CRP, IL-6, TNF-α, and MDA and decreased the serum levels of GSH and TAS (P<0.05) in diabetic rats. Oral administration of *Spirogyra* alleviated adverse effects of diabetes on oxidative stress and inflammatory factors in diabetic rats (P<0.05).

**Conclusion:** It can be stated that *Spirogyra* algae extract can be used for treatment of diabetes likely due to prevention of oxidative stress and alleviation of inflammation in the rat model.

## Introduction

Diabetes mellitus (DM) is a metabolic disorder and it is classified into two types, type 1 and type 2. Type 2 diabetes mellitus is broadly related with hyperglycemia and insulin resistance^1^. Insulin resistance in some tissues in the body, such as skeletal muscle, adipose tissue, and the liver, causes the development of diabetes^2^. It is reported that a positive correlation exists between levels of proinflammatory cytokines tumor necrosis factor-α (TNF-α), interleukin-1 (IL-1), and IL-6 with insulin^3^. An interesting study has suggested that inflammation can directly causes pathogenesis of insulin resistance^4^. TNF-α causes inflammation and decreases β-cell secretion of insulin^5^. Additionally, TNF-α increases triglycerides production^6^. On the other hand, oxidative stress is created due to an imbalance between oxidant and antioxidant systems, increase of free radical production, and reduction in activity of antioxidants^7^. Oxidative stress induces hyperglycemia, promotes diabetes, and causes other severe complications^8^. Antioxidants can scavenge free radicals and reactive oxygen species (ROS) through inhibiting lipid peroxidation and decreasing the adverse effects produced by ROS^9,10^. Generally, synthetic drugs are used to treat the diabetes, but their use is faced with major limitations due to their many side effects. Natural compounds have recently been tried in many regions with mixed results. In this context, algae are major source of natural compounds that are broadly found in nature. Algae are known to have antibacterial, antiviral, antioxidant, anti-inflammatory properties, and cytotoxic activities^11^. Polysaccharides in algae are also found to scavenge free radicals under *in vivo* and *in vitro* conditions^12^. *Spirogyra* spp. of algae are filamentous freshwater green algae that contain high amounts of protein, carbohydrate, fat, sulfate, and dietary fiber^13^. Moreover, the *Spirogyra* produce compounds such as alkaloids, steroids, flavonoids, tannins and terpanoids^14^ *Spirogyra neglecta* (SN) is reported to have antioxidant activity^15^ and anti-inflammatory effects^16^. Keeping in view the above mentioned studies, it was hypothesized that SN algae can be used to treat the diabetes patients due its known antioxidant and anti-inflammatory properties. Thus, this study was conducted to evaluate the effects of SN algae on oxidative stress and inflammation in diabetic the rat model.

## Materials and Methods

### Animals

Adult male Wistar rats, weighing 200±20 g and 9 wks of age, were purchased from Pastur Institute, Tehran-Iran. Sixty-four Wistar rats were randomly assigned into 8 groups (n=8). Animals were further divided into four groups for diabetic and four groups for non-diabetic. Animal in the diabetic groups received 0 (control), 15, 30, or 45 mg/kg of SN extract. Algae extract was administrated in gavage form for 42 days after induction of diabetes. Animals in healthy, nondiabetic groups orally received 0 (control), 15, 30 or 45 mg/kg of SN extract. Rats were grouped in a well-ventilated animal facility in stainless steel cages in order to allow them free mobility. Rats received a standard feed (Javaneh Khorasan, Iran) and fresh water. Rats were kept under controlled conditions including temperature (22 ± 2°C) and humidity (55 ± 5%). A lighting diet (12 h light/12 h dark) was considered. All the experiments were conducted in accordance with the National Institutes of Health Guide (NIH Publication No. 85-23, revised 1996) for the Care and Use of Laboratory Animals.

### Preparation of Spirogyra neglecta extract

*Spirogyra neglecta* was provided from Bistoon pond (Kermanshah-Iran) and dried for 10 d. The extract was prepared as described by Ontawong et al.^15^. Lyophilized extract were maintained in 4°C prior to experiments.

### Pilot study

Forty Wistar rats were fasted for 24 h and then divided into 8 groups with 5 animals each. Animals were orally administrated with 0.2 mL distilled water and a dose of SN extract at either 10, 15, 30, 45, 60, 70, or 90 mg/kg. The rats were observed in order to assess the behavioral responses, toxicity signs, and mortality for 48 h. We did not observe any mortality but diarrhea was observed in the 70 and 90 mg/ kg test groups. Thus, we selected doses of 15, 30, and 45 mg/kg for further studies. These doses were not observed to have any behavioral responses and signs of toxicity.

### Induction of Diabetes

In order to induce diabetes, animals were intraperitoneally administrated with 55 mg/kg of streptozotocin (STZ, Sigma, St. Louis, MA, USA) in 0.1 M citrate buffer pH 4.5 as described by Jaiswal et al.^17^. Rats were bled through the tail in order to confirm induction of diabetes, and assessed by a glucometer (Accu-Check, Roche, Manheim, Germany). Rats with a glucose level >250 mg/dl were considered diabetic^17^.

### Blood sampling and antioxidant, inflammatory and blood variables

On day 42 after interventions and 24 h fasting, animals were intraperitoneally anaesthetized by administration of sodium pentobarbital injection (60 mg/kg). Blood samples were collected into tubes without heparin. The obtained samples were centrifuged at 4000 g for 10 min in 4°C and sera were stored in −20°C for subsequent trials. An enzyme-linked immunosorbent assay (ELISA) was used to evaluate the levels of TNF-α and IL-6 as recommended by the manufacturer’s instructions. In short, rat TNFα and IL-6 specific-specific monoclonal antibodies were loaded into 96-well plates. The rat specific detection polyclonal antibodies were biotinylated. The samples test and biotinylated detection antibodies were added to the wells subsequently and then continued through washing with PBS or TBS buffer. Avidin-Biotin-Peroxidase Complex was included and unbound conjugates were investigated away with PBS or TBS buffer. TMB catalyzed by HRP used to produce a blue color product which altered into yellow in following inclusion of acidic stop solution. The density of yellow was proportional rather than TNFα level of sample captured in plate. TBARS (thiobarbituric acid reactive substances) procedure was used to evaluate the serum malondialdehyde (MDA) as described by others^18^. The levels of glutathione (GSH) was evaluated by Ellman’s reagent (5, 50-dithio-bis-2-nitrobenzoic acid,) as reported by previous studies^19^. The total antioxidant status (TAS) was assessed by FRAP (ferric reducing antioxidant power) procedure^20^. C-reactive protein (CRP) was measured by specific kits (The Invitrogen CRP ELISA Kit, Camarillo, CA). The serum concentrations of glucose, cholesterol, triglycerides, and insulin were assessed by commercial kits (Pars Azmoon-Iran).

### Statistical analysis

The data were analyzed by using GraphPad Prism statistical software (GraphPad Software, Inc., La Jolla, CA, USA) by one-way/two way analysis of variance (ANOVA) followed by Dunnett’s Multiple Range Test. Animals in healthy group and diabetic group were separately compared by T-test. The findings are shown as mean ± SD. A value of *P*<0.05 was considered to be statistically significant.

## Results

### Antioxidant variables

Effects of different levels of SN extract on MDA, GSH, and TAS are shown in Figure 1. Results showed that induction of diabetes significantly increased levels of MDA (Figure 1A), and decreased levels of GSH and TAS (*P* < 0.0001) (Figure 1B and C, control diabetic vs. control non-diabetic). The different doses of SN had no significant effects on MDA, GSH, and TAS in non-diabetic groups (*P* > 0.05). In diabetic rats, with increasing doses of extract, serum levels of MDA decreased and levels of GSH and TAS increased (*P* < 0.0001). The best responses were observed in 15, 30, and 45 mg/kg of SN extract in diabetic group, respectively.

**Figure 1.**
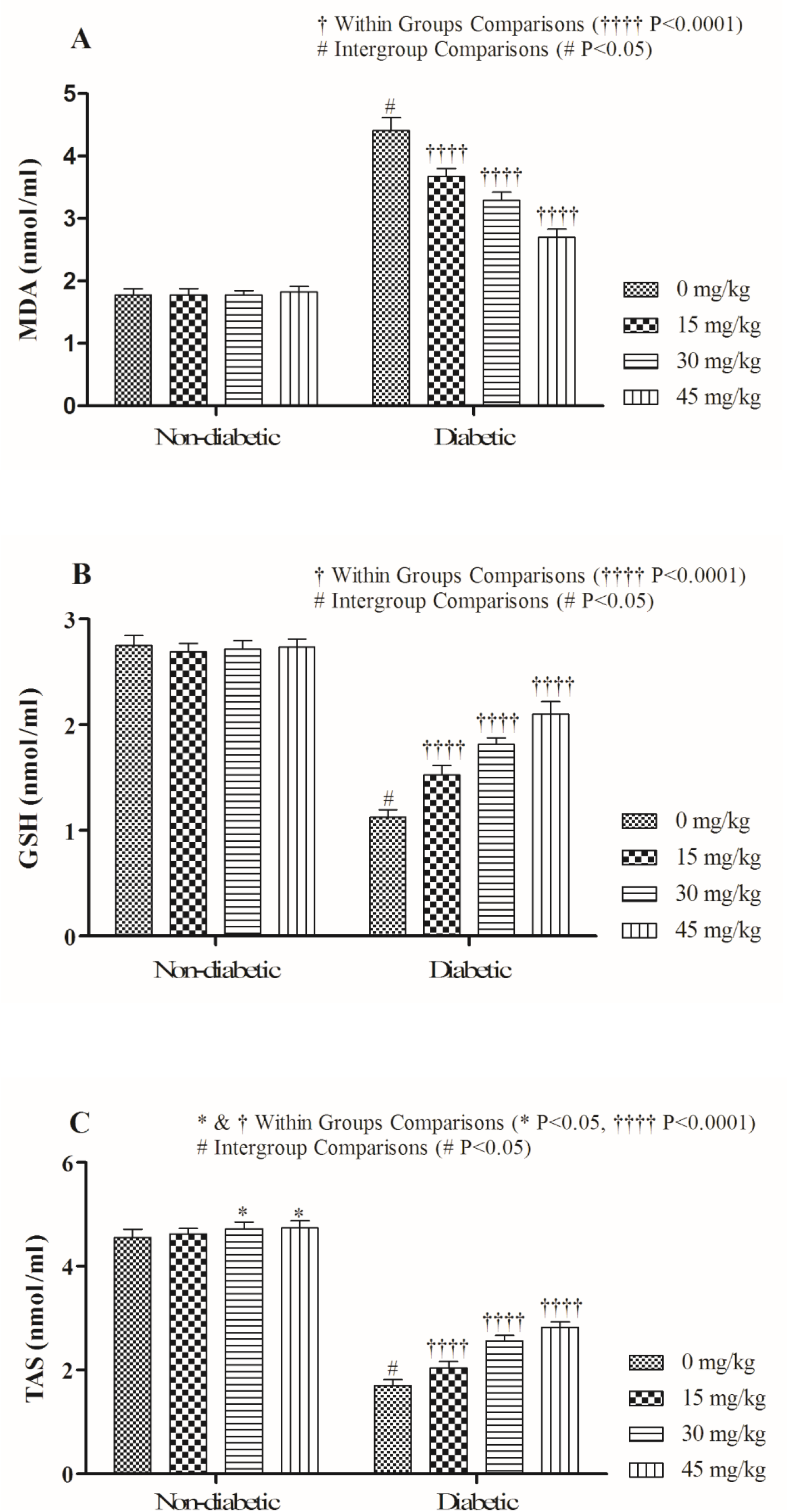
Effects of different levels of SN extract on antioxidant variables in diabetic and non-diabetic rats

### Inflammatory variables

The serum concentrations of TNF-α, IL-6, and CRP were significantly increased in control diabetic in comparison to non-diabetic groups (Figure 2, *P* < 0.0001). Oral administration of SN extract especially in higher doses reduced the levels of TNF-α, IL-6, and CRP.

**Figure 2.**
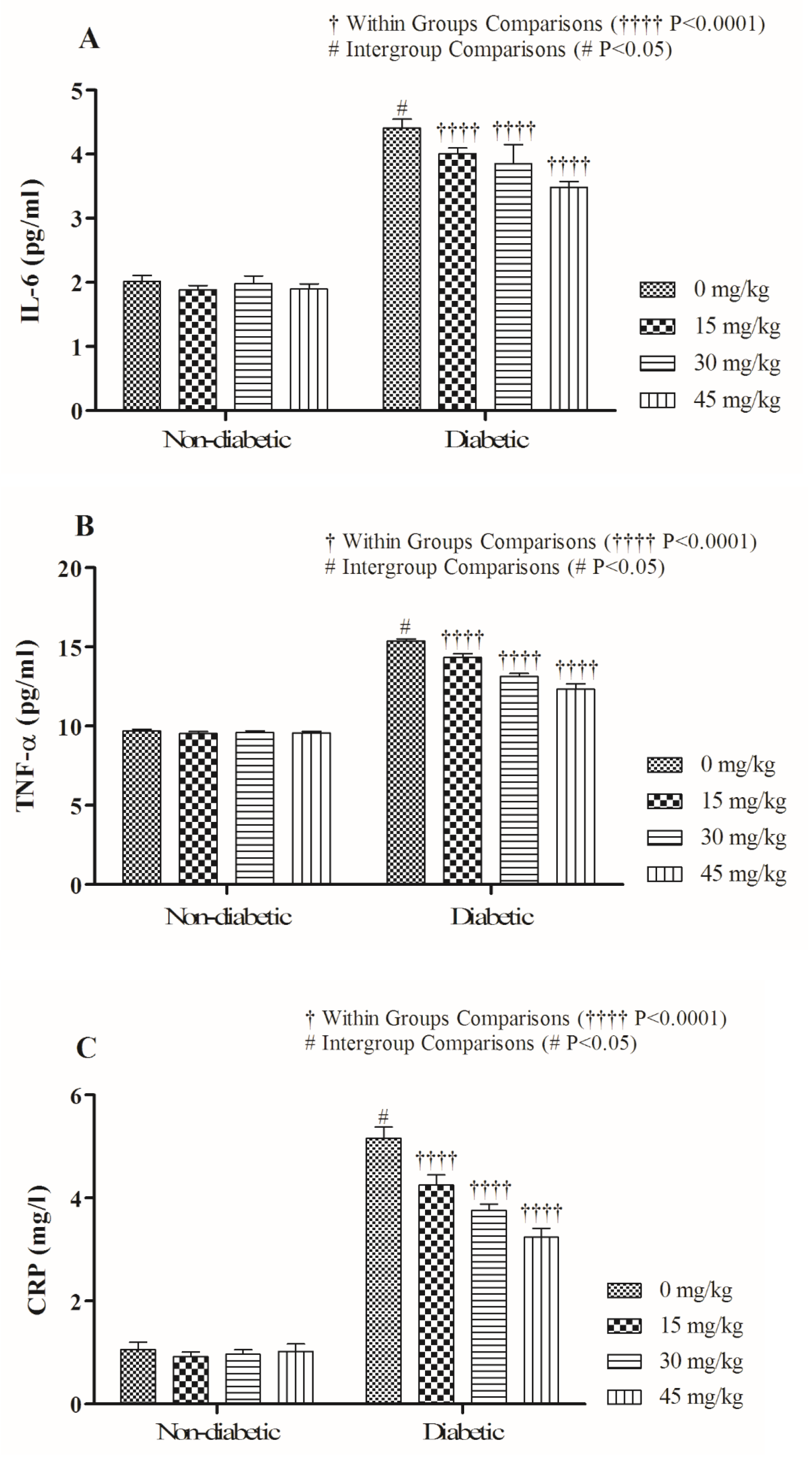
Effects of different levels of SN extract on inflammatory variables in diabetic and non-diabetic rats

### Blood variables

The serum concentrations of cholesterol, triglycerides, and glucose were significantly increased and insulin concentration was decreased in diabetic rats in comparison to non-diabetic rats (Figure 3, *P* < 0.0001). Diabetic rats treated with SN extract showed lower levels for cholesterol, triglycerides, and glucose and higher levels for insulin in comparison to diabetic control (*P* < 0.0001). Better responses were observed in diabetic rats treated with higher doses (30 and 45 mg/kg).

**Figure 3.**
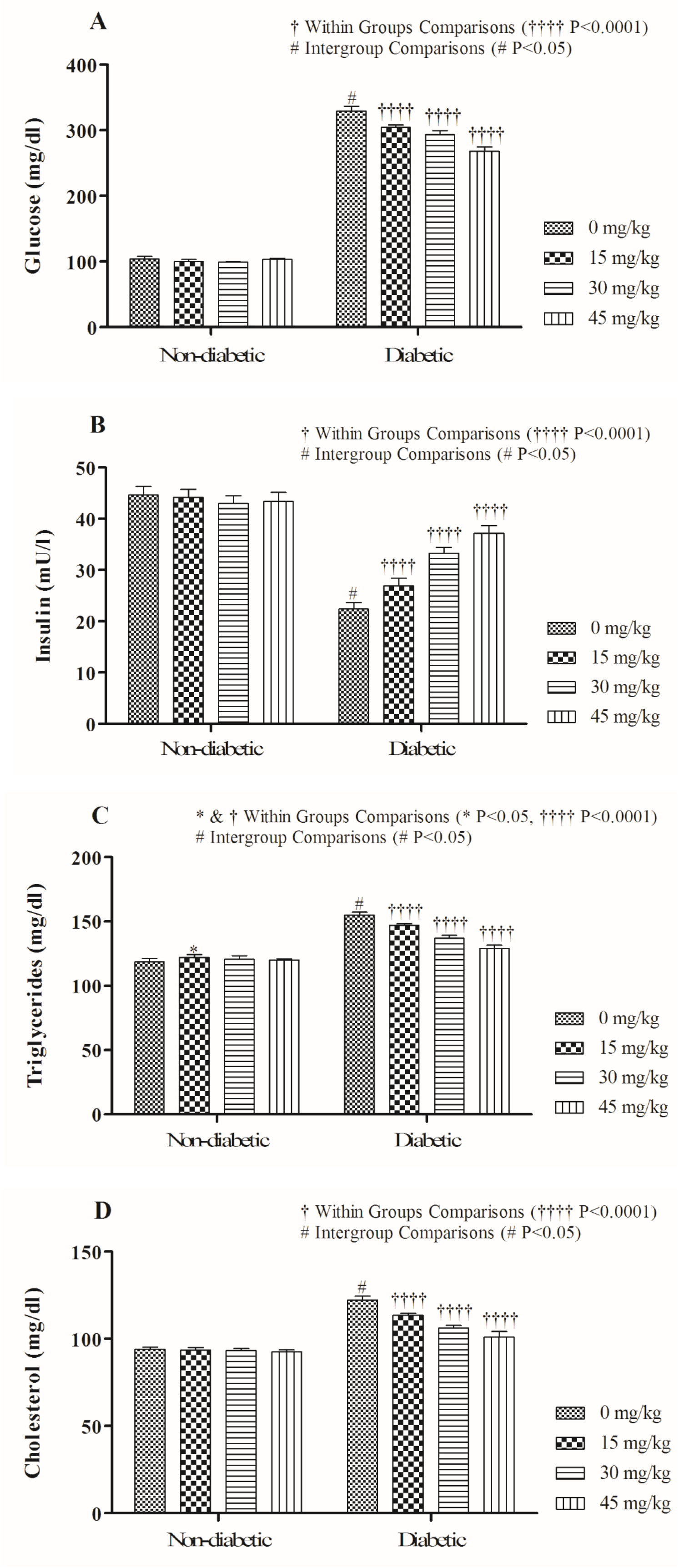
Effects of different levels of SN extract on blood variables in diabetic and non-diabetic rats

## Discussion

In this study, we have evaluated the pro-oxidant-antioxidant balance by assessing MDA, GSH, and TAS levels in the diabetic and non-diabetic rats. Comparison of control diabetic and healthy rats showed a balance change toward pro-oxidation in STZ-induced diabetic animals. This resultantly increased the MDA, and decreased the GSH and TAS in rats.

Previous published studies have shown that STZ induces diabetes and increases susceptibility to lipid peroxidation^7,21^. This might explain the phenomena behind the oxidative stresses induced diabetes and pathogenesis of diabetes. In this relation, the MDA is a product of lipid peroxidation that is rapidly combined with biomolecules and deregulates glucose metabolism^22^. In the present study, oral administration of SN extract in diabetic rats improved antioxidant status due to increased GSH and TAS, and decreased MDA. This could be attributed to antioxidant components found in SN extract. SN extract is well known to have antioxidant properties due to its polyphenols contents^23,24^. In a relative study, Ontawong et al.^15^ showed that SN extract can directly scavenge free radicals and, thus, decrease MDA. It is well known in the literature that improved antioxidant status decreases lipid peroxidation and MDA levels. It can be speculated that SN extract is increasing the TAS, and ultimately the GSH could decrease levels of MDA.

On othe hand, the serum concentrations of TNF-α, IL-6, and CRP were significantly increased in diabetic rats, but oral administration of SN extract especially in higher doses reduced the levels of TNF-α, IL-6, and CRP. Oxidative stress causes production of abnormal cytokine production (TNF-α and IL-6)^25^. TNF-α not only increases adipocyte lipolysis but also has adverse effects on the insulin signaling pathway through changing the tyrosine/serine phosphorylation of insulin receptor substrate^26^. CRP is a key inflammatory factor that is regulated by IL-6, IL-1, and TNF-α, and produced by the liver in response to inflammation^27^.

In relation to this, Virgolici et al.^28^ showed that increased levels of inflammation were related with the increased oxidative stress. We also observed and noticed that diabetes induces oxidative stress in rats. In addition, Chang et al.^29^ have reported that diabetes mellitus may induce nuclear factor kappa B (NF-κB) that is responsible for the production of some proinflammatory cytokines. In our study, the oral treatment of SN extract interestingly decreased the levels of inflammatory cytokines. Similarly, Nasirian et al.^30^ reported that the oral gavage of *Spirulina platensis* microalgae reduced inflammatory factors in diabetic rats. Improved inflammatory parameters in response to SN extract can be attributed to antioxidant activity of polyphenolic compounds existing in SN extract that prevents the development of ROS. As mentioned earlier, SN extract has antioxidant activity that may prevent production of inflammatory cytokines. Nevertheless, greater antioxidant activity was observed in higher levels of its doses in the rat model.

Hypertriglyceridemia and hypercholesterolemia are common signs in diabetes^31^. Previous studies have also shown that STZ increases the sensitivity to lipid peroxidation^7,21^. In addition, this might be the phenomena in which the STZ enters into pancreatic β cell through the low-affinity glucose protein-2 transporter. The STZ damages insulin-producing islet β-cells and decreases insulin secretion^7^. In this complex pathway system, the reduced hepatic insulin sensitivity may be caused by an increase in hepatic gluconeogenesis, postprandial hyperinsulinemia, and increased production of triglycerides in the liver cells. SN extract has alleviated hyperglycemia, dyslipidemia and increased the insulin sensitivity in the rat model with diabetes mellitus^33^. Thus, TNF-α over produces triglycerides in plasma^6^. TNF-α reduces glucose intake of peripheral tissues through targeting insulin signaling pathways and glucose transporter 4 (GLUT4)^32^. In the present study, oral supplementing of SN extract have decreased the levels of TNF-α. This might improve the levels of insulin and glucose through decreasing TNF-α. Moreover, the oxidative stress and reactive oxygen species also degrades the lipids and increases the lipid peroxidation and ultimately leads to lipolysis. It can be concluded that SN extract decreased the levels of lipids probably by antioxidant properties in the rat model.

## Conclusion

Our overall findings showed that STZ increased inflammatory and lipid variables in diabetic rat model. While decreasing antioxidant variables and levels of insulin. Oral treatment of SN extract has not cause any significant changes in the inflammatory, antioxidant, insulin, or lipid variables in non-diabetic rats. Oral gavage of SN extracts could decrease levels of MDA, lipid peroxidation and inflammatory variables in diabetic rats. It could be concluded that the SN extracts improved blood biochemical variables probably by acting on antioxidant pathway system and thus reducing inflammatory variables. Since levels of 45 mg/kg of SN extract in the rat model have better effects, we therefore recommend the same or slightly higher levels for treatment and management of diabetes in future trials.

## Author contributions

All authors contributed toward data analysis, drafting and revising the paper and agree to be accountable for all aspects of the work.

